# Highly parallel genomic selection response in replicated *Drosophila melanogaster* populations with reduced genetic variation

**DOI:** 10.1101/2021.04.06.438598

**Authors:** Burny Claire, Nolte Viola, Dolezal Marlies, Schlötterer Christian

**Author notes:** equal contribution (Burny Claire, Nolte Viola). Corresponding author: Christian Schlötterer.

## Abstract

Many adaptive traits are polygenic and frequently more loci contributing to the phenotype than needed are segregating in populations to express a phenotypic optimum. Experimental evolution provides a powerful approach to study polygenic adaptation using replicated populations adapting to a new controlled environment. Since genetic redundancy often results in non-parallel selection responses among replicates, we propose a modified Evolve and Resequencing (E&R) design that maximizes the similarity among replicates. Rather than starting from many founders, we only use two inbred *Drosophila melanogaster* strains and expose them to a very extreme, hot temperature environment (29°C). After 20 generations, we detect many genomic regions with a strong, highly parallel selection response in 10 evolved replicates. The X chromosome has a more pronounced selection response than the autosomes, which may be attributed to dominance effects. Furthermore, we find that the median selection coefficient for all chromosomes is higher in our two-genotype experiment than in classic E&R studies. Since two random genomes harbor sufficient variation for adaptive responses, we propose that this approach is particularly well-suited for the analysis of polygenic adaptation.

## INTRODUCTION

Many adaptive traits have a polygenic basis (Barton and Etheridge, 2018; Barghi et al, 2019; Hoffman et al, 2003), where typically more contributing loci are segregating in a population than needed to reach the trait optimum (Yeaman, 2015). For highly polygenic traits, the contribution of a single locus during adaptation to a new environment, *i*.*e*. a new phenotypic optimum, will be small, usually too small to be detected by classic population genetic tests (Pritchard et al, 2010; Pritchard and Di Rienzo, 2010). Thus, tests for polygenic adaptation aggregate signals across multiple loci to gain statistical power (Turchin et al, 2012; Berg and Coop, 2014; Sella and Barton, 2019). However, distinguishing the contributions of demography and selection in these aggregated signals can be challenging in natural populations because of residual population structure (Barton et al, 2019; Sohail et al, 2019; Berg et al, 2019). Hence, experimental evolution has been proposed as an alternative approach to study polygenic adaptation (Barghi et al, 2020; Lou et al, 2020; Vlachos and Kofler, 2019). Laboratory natural selection within the Evolve and Re-sequencing (E&R) framework (Garland and Rose, 2009; Turner et al, 2011; Long et al, 2015; Schlötterer et al, 2015) has been successfully used to study adaptation in controlled environments, combining experimental evolution and Pool-sequencing on replicated populations (Schlötterer et al, 2014).

Simulation studies (Baldwin-Brown et al, 2014; Kofler and Schlötterer, 2014; Kessner and Novembre, 2015) recommend optimizing different design parameters to obtain a good mapping resolution. An established strategy is to use a large number of founder genotypes. Maximizing the number of founders provides the advantage that the contributing alleles segregating at intermediate frequency will be located on multiple haplotypes, which facilitates their identification (e.g. Kelly and Hughes, 2019). However, for highly polygenic traits, increasing the number of founders also increases the number of available contributing alleles, which may either trigger competition between the present haplotypes if they interfere with each other (Hill and Robertson, 1968), or inflate genotypic redundancy making evolution less repeatable (Láruson et al, 2020). Additionally, increasing the number of founders lowers their starting frequency, which in turn increases their chance to be lost by drift.

As a consequence, a (highly) heterogeneous response between replicates is expected and has been seen in several E&R studies (e.g. Seabra et al, 2017; Griffin et al, 2017; Hardy et al, 2018; Barghi et al, 2019; Rêgo et al, 2019) – even when the same founder population is used, and in particular in small populations where stochastic sampling effects have a strong impact on allele frequencies. Nevertheless, various E&R studies displayed (highly) parallel selection signatures despite using a large number of founders (e.g. Martins et al, 2014; Burke et al, 2014; Graves et al, 2017; Phillips et al, 2018; Kelly and Hughes, 2019). These conflicting observations imply that our understanding of the adaptive response, *i*.*e*. the adaptive architecture, in E&R studies is not yet complete and more data are required to evaluate which factors contribute to (non-)parallel selection responses (Barghi and Schlötterer, 2020; Otte et al, 2020; Matos et al, 2015).

Parallel genomic responses are a key factor determining the power of many statistical tests to detect selection at a given locus in E&R studies (reviewed in Vlachos et al, 2019). The degree of parallelism depends on the probability that a particular favorable allele from standing genetic variation will respond to selection, with loci of large effect showing more parallel signatures (Hermisson and Pennings, 2017; Castro et al, 2019). Various experimental design parameters determine how concordant the selection responses are (Vlachos and Kofler, 2019; Baldwin-Brown et al, 2014; Kofler and Schlötterer, 2014; Kessner and Novembre, 2015). First, higher starting frequencies and larger population size reduce the probability of stochastic loss and help to consistently detect not only large-effect but also moderate-effect loci. Second, depending on the distance to the new phenotypic optimum, either more sweep-like (distant) or shift-like (less distant) responses are favored (Matuszewski et al, 2015; Christodoulaki et al, 2019; Hayward and Sella, 2019). Third, with increasing redundancy, the selection response is becoming less parallel (Láruson et al, 2020).

In this study, we designed an experiment which aims to achieve a highly parallel selection response across replicates by accounting for all three factors outlined above. Given that many adaptive variants are present in natural *Drosophila* populations, we drastically reduced the amount of segregating variation in the founder population by using only two founder genotypes. We first created 10 replicate populations from two parental inbred *D. melanogaster* strains, Samarkand and Oregon-R. We then exposed the replicate populations to an extreme temperature regime (constant 29°C), which is only slightly below the maximum temperature at which *D. melanogaster* are viable and fertile (Fig 1, Hoffmann, 2010). Eventually, all contributing alleles that start at intermediate frequency in the founder population will be measured after 20 generations.

**Figure 1.**
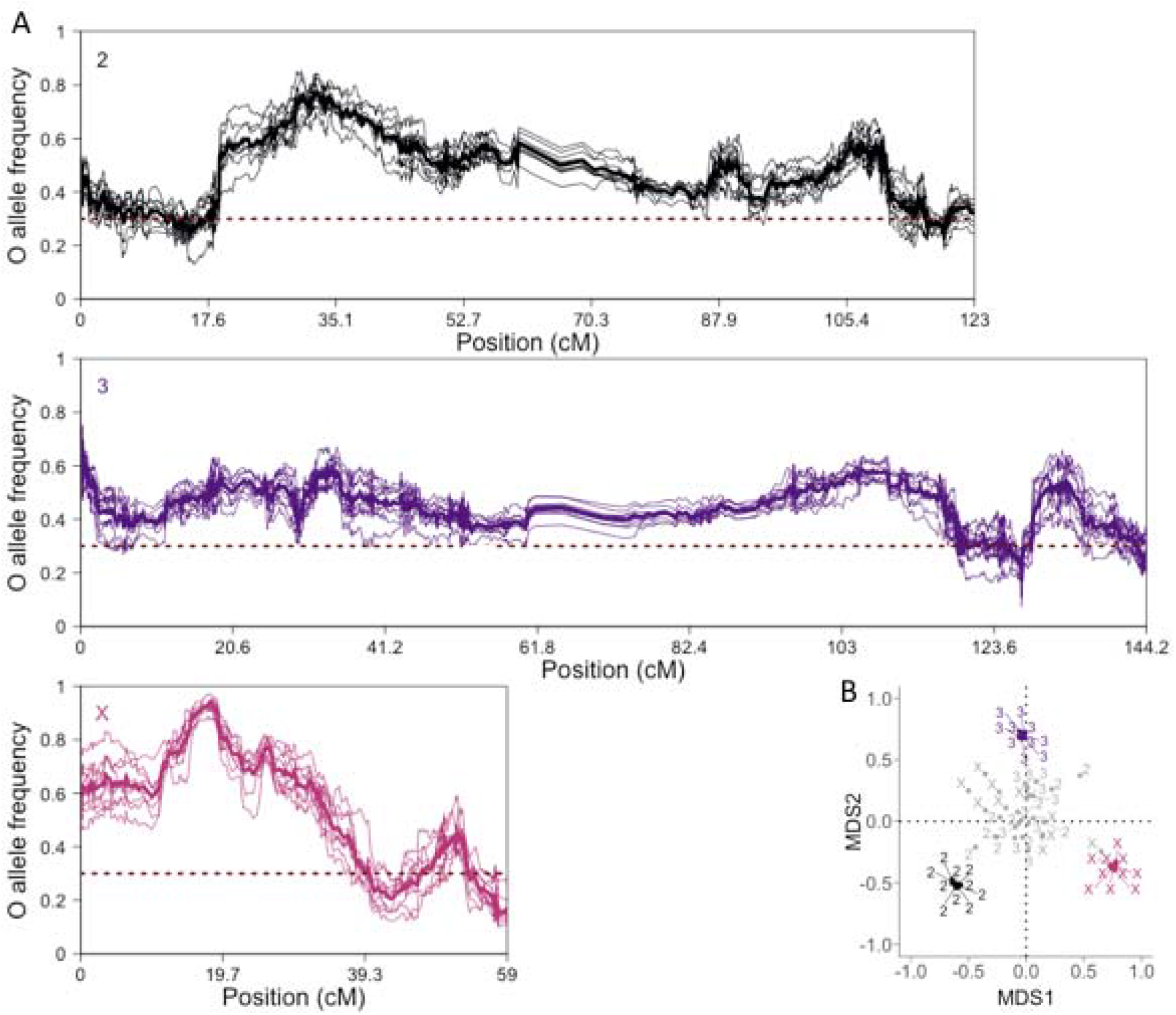
Strong parallel response after 20 generations of evolution at 29°C. A) Smoothed Oregon-R (O) allele frequency (y-axis) at F20 in all replicates colored by major chromosome in cM unit (x-axis). The same color code applies to all figures (from dark to light: chromosomes 2, 3 and X). The median O allele frequency (AF) is computed over non-overlapping windows of 250 SNPs. The bold line represents the median O AF per window over the 10 replicates and the horizontal dotted line the starting O AF (0.3). B) 2D Multidimensional Scaling (MDS) projection of the pairwise ρ Spearman correlation matrix between empirical (colored) and neutral (gray) allele frequencies per major chromosome. The correlation coefficient values were transformed to distances (2√(1-ρ)) prior to projection.

By analyzing the genomic responses in the 10 replicate populations maintained for 20 generations at a hot temperature, we find that two founder genotypes harbor enough natural variation to ensure a selective response. A very strong and highly parallel selection signature is seen in all replicates. This demonstrates that even for temperature adaptation, which is highly polygenic, an adequate experimental design, *i*.*e*. a reduced founder diversity and a distant trait optimum, results in reproducible selection signals.

## RESULTS

### Parallel response after 20 generations of evolution at high temperature

Using two genotypes to set up the founder population provides the advantage that all parental alleles start from the same frequency across the entire genome. A simple genome-wide allele frequency plot along the genome provides an intuitive visualization of the selection targets (Fig 1A): the pronounced allele frequency increase of the putatively selected alleles, either Oregon-R (AF>30%) or Samarkand (AF<30%), generates a “hill-valley-like” landscape. Since recombination rate *a priori* determines the width of the genomic region affected by a selected site (Felsenstein, 1974; Barton, 1995; Otto and Lenormand, 2002; Roze and Barton, 2006), we scaled the chromosomes in cM unit (for a base-pair scaling, see Fig SI 4). Throughout the entire genome, we observe a fast and strong response after 20 generations (Fig 1A) where in all replicates, large, linked genomic regions experience very similar changes in frequency.

The high level of parallelism among the empirical replicates is reflected in highly correlated allele frequencies between replicates, higher than 0.8 (t-test on pairwise Spearman correlation coefficient ρ per arm; mean ρ_2_=0.89 (t(40)=200, adjusted (adj.) p<1.7×10^−65^), mean ρ_3_=0.80 (t(40)=100, adj. p<6.4×10^−55^), mean ρ_X_=0.92 (t(40)=200, adj. p<3.9×10^−67^)). Such high correlations are not observed among replicate populations in neutral simulations (t-test on pairwise ρ per arm; mean ρ_2_=0.04 (t(40)=0.8, adj. p>0.57), mean ρ_3_=0.07 (t(40)=2, adj. p>0.37), mean ρ_X_=0.0 (t(40)=0.09, adj. p>0.95)). We visualized the difference between the empirical and simulated replicates by projecting the pairwise correlation matrix in a two-dimensional multidimensional scaling plot (Fig 1B), which highlights the similarity between the empirical replicates for each major arm, whereas in the neutral simulations no clustering of replicates was apparent.

While it is difficult to provide a statistically sound estimate of the number of selection targets, it is apparent that reducing the genetic variation to two genotypes still leaves a considerable reservoir of favorably selected alleles. This strong selection response is also reflected in effective population size (*N*_*e*_) estimates based on allele frequency changes. For the X chromosome, *N*_*e*_ barely reaches 25 with a median of 21 and is also rather small on the autosomes (median of 55, SI Table 2), given a census size of 1,500 flies in each replicate. The effective population size on the X chromosome is much lower than the expected 3/4 reduction relative to the autosomes (Charlesworth, 2009). This implies that the efficacy of selection differed between the autosomes and X and that selection was considerably stronger on the X chromosome (see discussion for possible explanations).

The experiment started from two genotypes and in 20 generations the number of recombination events that can uncouple contiguous blocks of Oregon-R/Samarkand alleles which experience a strong frequency increase is limited. This non-independence of neighboring sites translates into the “hilly” landscape of allele frequency changes. In the absence of haplotype data from the evolved flies, we used the loss of autocorrelation in allele frequency as a proxy for the decay of linkage disequilibrium to quantify the association between genomic sites (Fig 2A). The correlation between increasingly distant windows decayed faster on the autosomes (with a median of 5.9Mb and 4.7Mb over the 10 replicates for chromosomes 2 and 3) compared to the X chromosome (median of 6.6Mb) (Fig 2B), implying less LD on the autosomes. We attribute the independence of neighboring windows at a lower distance on the autosomes (correlation outside 95% confidence interval) to differences in selection intensities: stronger selection reduces the effective population sizes beyond the 3/4 expected from the ratio of X chromosomes to autosomes, which results in less opportunity for recombination on the X chromosome.

**Figure 2.**
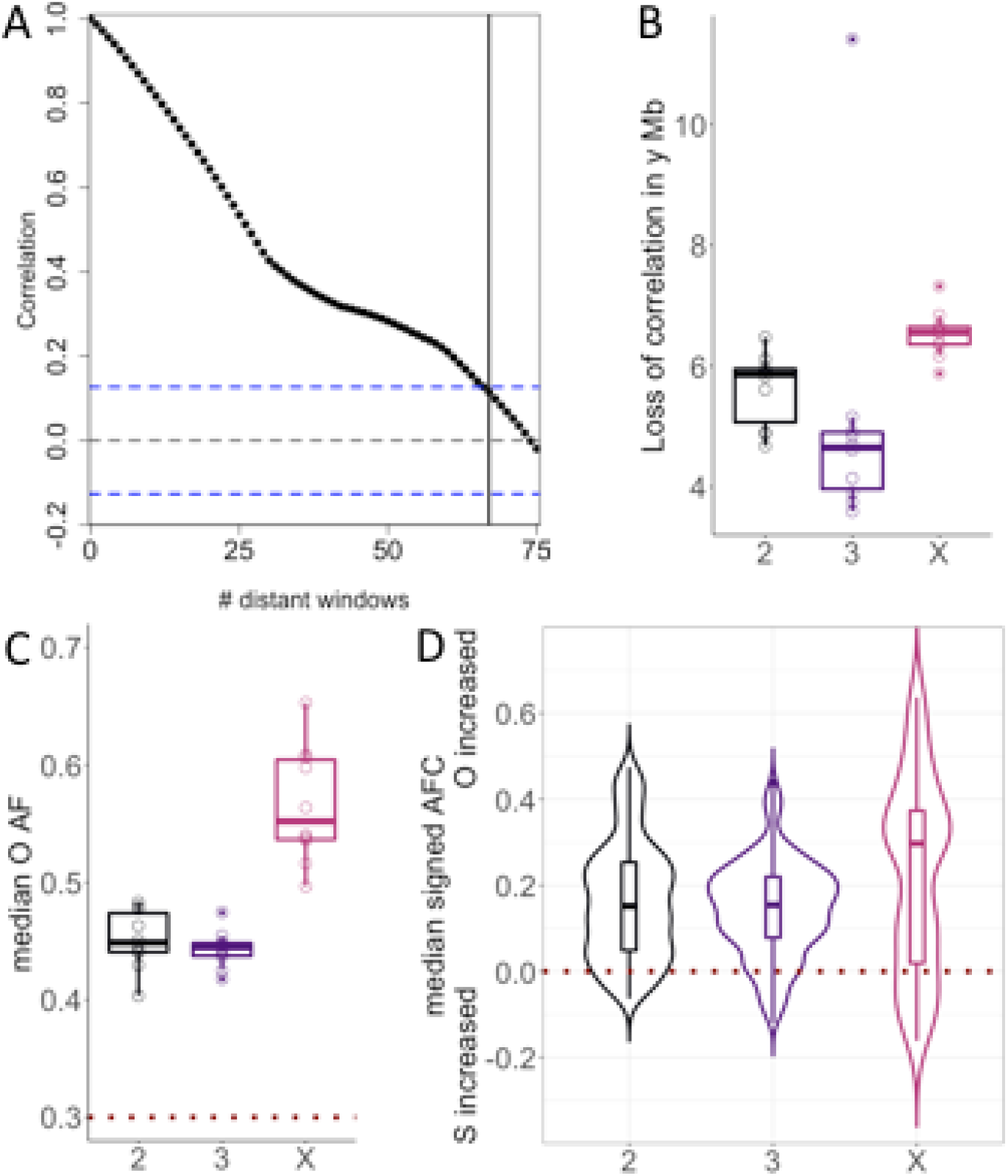
Quantification of the evolutionary response at F20. A) and B) Loss of correlation at the major chromosomes. A) Example of a scatterplot of ρ Spearman correlation against distance between two windows measured in the number of windows separating them. The blue dotted lines represents +/-1.96/√m, with m number of windows. B) Jittered boxplots of physical distance in Mb where Linkage Equilibrium (LE) is reached at a 5% threshold (vertical black line in the A panel). C) Jittered boxplots of median O allele frequency (AF) on the major chromosomes in each replicate. D) Boxplots overlaid with violin plots of AFC. A positive (negative) allele frequency change (AFC) indicates that the O genotype increases (decreases) in the window relative to the starting frequency of 0.3. The horizontal dark red dashed line indicates no change in frequency after 20 generations.

At 29°C the two separated parental lines suffered similarly from the high temperature regime and produced low numbers of offspring (data not shown). When the two strains were combined in the experimental evolution cage, the Oregon-R alleles clearly outcompeted the Samarkand genotypes (Fig 1A, Fig 2C,D): the median Oregon-R AFC was significantly higher than 0 (0.15, 0.15, 0.30 for chromosomes 2, 3 and X; adj. p<3.5×10^−89^, adj. p<7.7×10^−110^, adj. p<4.6×10^−18^ on each sign test; Fig 2D). Although some heterogeneity can be observed along the chromosome arms (Fig 1A; median coefficient of variation is 0.10, 0.11, 0.14 for chromosomes 2, 3 and X), the median Oregon-R allele frequency increased on each chromosome, ranging from 40% to 65% (Fig 2C), which suggests a genome-wide rather than an isolated footprint of selection.

### Exceptionally strong, genome-wide selection signatures

With all alleles occurring at similar frequency throughout the entire genome, the comparison of allele frequency changes provides a direct readout of the selective force operating on each SNP - either directly or through linkage to selection targets. To compare the selection experienced in this two-genotype experiment to two other short-term *Drosophila* E&R studies (Table 1) that differ in the number of founders (>200) and consequently in the distribution of starting allele frequencies, we transformed the allele frequency changes into selection coefficients, *s*, which allows the comparison of alleles with different starting frequencies. The pronounced differences in median absolute *s* between the X chromosome and autosomes were specific to the two-genotype experiment (Fig 3, estimates on x-axis). Across all chromosomes the median absolute *s* was significantly higher for this study compared to the two other studies (Fig 3, estimates on x-axis and adj. p). This clearly indicates that the two E&R studies experienced less selection, not only on the X chromosome, but genome-wide which may reflect the lower temperature (23°C and 25°C) during their maintenance. The differences in selection intensity between the two-genotype experiment and E&R studies with many founder genotypes are also reflected in effective population size (*N*_*e*_) estimates. With *N*_*e*_ estimates not higher than 60 and 26 for the autosomes and X in all replicates (SI Table 3), *N*_*e*_ of this study was considerably lower than for the two other E&R studies (see Fig 3 legend), suggesting that a much larger fraction of the genome experienced drastic allele frequency changes.

**Table 1.** E&R datasets information.

| Number founders | Census size | Pressure | Species | Generation picked | Sequencing information | Publication |
| --- | --- | --- | --- | --- | --- | --- |
| 2 | 1,500 | LNS<br>constant 29°C | *D. melanogaster* | 20<br>non<br>overlapping | Pool-seq of<br>1500 mixed<br>males and<br>females | this study |
| 202 | 1,000 | LNS<br>Fluctuating<br>temperature<br>(28°C/18°C,<br>mean 23°C) | *D. simulans* | 20<br>non<br>overlapping | Pool-seq of<br>1000 mixed<br>males and<br>females | Barghi et al,<br>2019 |
| 500 | mean = 1,187<br>range = 963–1,620<br>for the used<br>replicate | LNS<br>Constant<br>temperature<br>(25°C) | *D. simulans* | 15<br>overlapping | Pool-seq of<br>500 males and<br>females | Kelly and<br>Hughes, 2019 |

**Figure 3.**
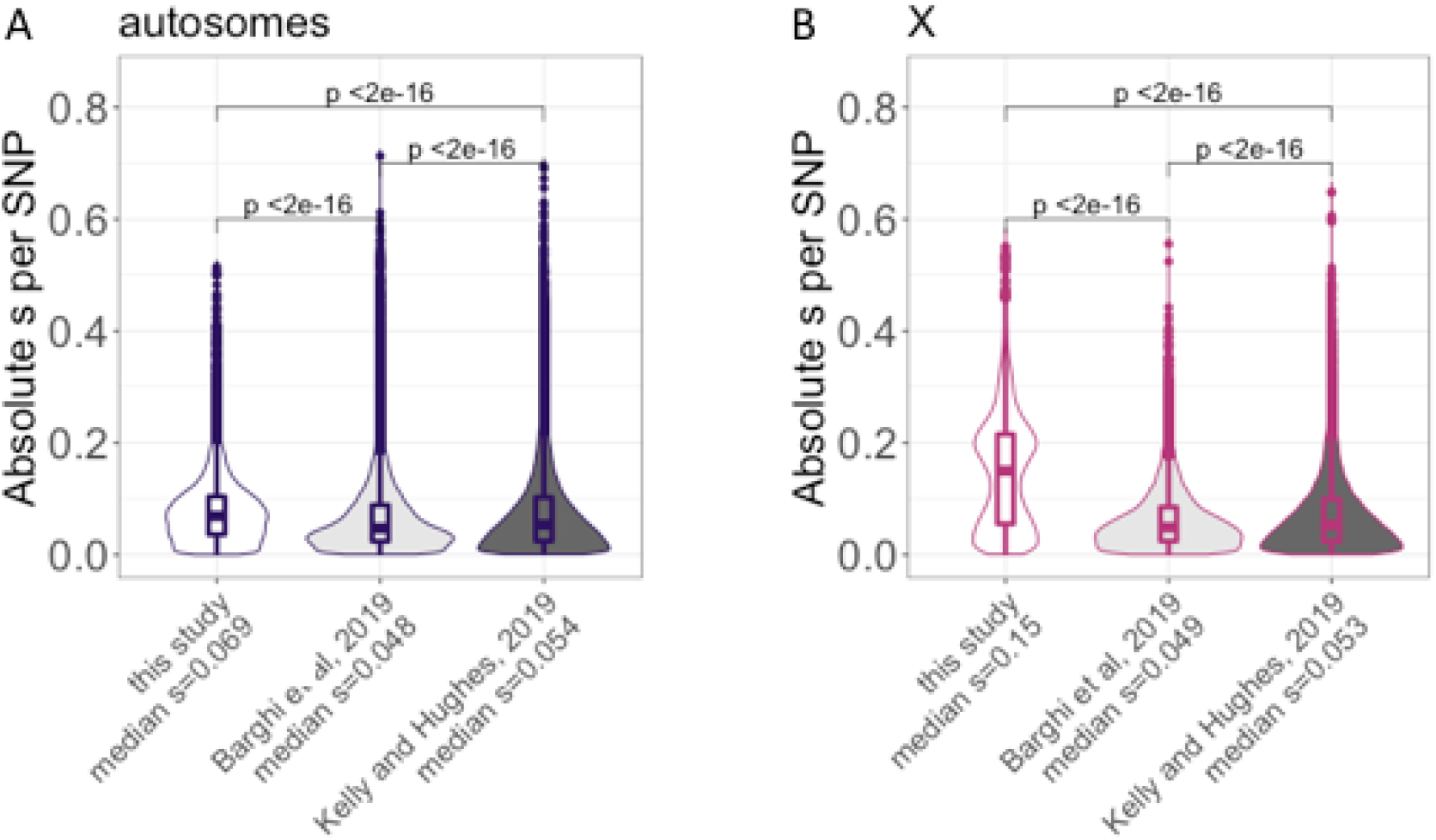
Distribution of absolute selection coefficients *s* per SNP across empirical E&R studies for the autosomes (A, 248,886 SNPs) and the X chromosome (B, 42,386 SNPs). Boxplots overlaid with violin plots per study (x-axis) where the median absolute *s* is reported. Adjusted p-values from pairwise Wilcoxon tests are indicating. Population size estimates of the autosomes (X) were 55 (14), 243 (227), 393 (359) for the three studies.

## DISCUSSION

The idea to start an experimental evolution study with only two genotypes is radically different from current E&R designs, but has already been used before in experimental evolution (Barnes, 1968; Kearsey and Barnes, 1970; Nuzhdin et al, 1998). While strong responses were observed, the link between genotypes and phenotypes the small population sizes used for example in the mouse selection experiments can result in considerable genetic heterogeneity among replicates which limits the power to detect loci with small/moderate effects. An interesting modification of the two genotype design has been used in fruit flies. From a polymorphic population two haplotype classes were identified with moderate number of linked allozyme markers, but each haplotype class harbored considerable variation which was not surveyed (Clegg et al, 1976). Evolving populations founded by these two haplotype classes showed very strong selection signatures, but the genomic response between the replicates was heterogeneous, which was attributed to genetic heterogeneity at the unmonitored part of the haplotype classes (Clegg et al, 1976). Overall, previous two genotype experimental evolution studies were primarily designed to study the phenotypic response, but not to obtain highly parallel genomic selection signatures among replicate populations.

In contrast, this study obtained a highly parallel selection signature which can be attributed to the use of two high frequency genotypes in the founder populations in combination with large census sizes (1,500 individuals). Such a highly parallel selection response provides an excellent tool to study adaptation because selection responses can be reliably distinguished from stochastic patterns - even with a small number of replicates. We propose that two-haplotype E&R studies can be used to experimentally confirm candidate alleles that were previously identified - similar to a secondary E&R experiment (Burny et al, 2020). One further advantage of the highly parallel selection signature seen in this two-haplotype E&R study is that it offers the opportunity to explore epistatic interactions when only a small number of loci are selected. Crossing one inbred strain to at least two other inbred strains (in separate pairwise crosses) provides an excellent system to study epistasis by contrasting the selection response of a candidate locus in different genomic backgrounds. The highly parallel response provides sufficient power to detect even small differences, *i*.*e*. changes in frequency of the same selection target, due to the genetic background.

For a highly polygenic architecture the selection response of a two haplotype E&R reflects the net effect of multiple contributing alleles in a selected haplotype block. A similar scenario has been modelled where an admixed genotype is broken up into haplotype blocks, which could introgress when the net effect of all loci in the haplotype block was positive (Sachdeva and Barton, 2018). If our two-genotype experiment is extended for more generations, the high parallelism of this set up can be used to study the breaking haplotype blocks of contributing alleles by stochastic recombination. This has been done in a recent E&R study in budding yeast, which also started from two inbred founder genotypes, but with a much larger population size and for 960 generations (Koshelava and Desai, 2018). Consistent with a highly polygenic architecture, the fitness of sexual populations continuously increased throughout the entire experiment, possibly by the creation of favorable allelic combinations during the experiment (Hickey and Golding, 2018). More generations are needed for the *Drosophila* experiment to determine whether fitness continues to increase as in the yeast study or plateaus when the trait optimum is reached (Franssen et al, 2017, Höllinger et al, 2019).

Strong selection responses in populations derived from two founder genotypes imply that one allele provides an advantage relative to the other. While it is tempting to speculate that the fitness advantage is related to the temperature stress imposed during the experiment, we cannot rule out that the selection response is caused by a deleterious allele that was acquired during the long-term maintenance, since Samarkand and Oregon-R isofemale lines have been collected more than 90 years ago (Lindsley and Grell, 1968). Isofemale lines are typically maintained at small population sizes, which renders most mutations effectively neutral (Ohta, 1973; Kimura, 1983) and could lead to the accumulation of deleterious alleles that are fixed in the parental strains. Consistent with the presence of deleterious alleles, we noticed that heterozygous F1 flies produced a larger number of eggs at 29°C than the inbred strains which had difficulties to sustain the next generation. If deleterious alleles are the primary driver of the observed allele frequency changes, the predominant increase of Oregon alleles would suggest that Samarkand has accumulated more deleterious alleles than Oregon. This conclusion is not supported by obvious fitness differences of the two parental genotypes at 29°C. Alternatively, the lack of clear fitness differences in the parental lines could be explained by overdominance, but the reason for the predominant frequency increase of Oregon allele frequencies remains unclear. Additional generations at 29°C would help to distinguish between both explanations. Deleterious alleles would be ultimately purged while overdominance would result in a stable equilibrium frequency. A third interpretation of the data is based on epistatic interactions between Samarkand and Oregon alleles. If a few Samarkand alleles interact with many Oregon alleles, this could account for the advantage of heterozygotes and the predominance of Oregon alleles among the selectively favored ones. Epistatic interactions could be further tested when the Oregon genotype is competed with other genotypes in separate pairwise competition experiments.

A particularly interesting result was the different selection signature on the X chromosome compared to the autosomes. More pronounced allele frequency changes, and hence higher selection coefficients, were found on the X chromosome translating in lower *N*_*e*_ estimate than expected, *i*.*e*. lower than ¾ of the *N*_*e*_ on the autosomes. We propose two not mutually exclusive explanations for this observation: 1) the selected loci may be (partially) recessive which allows for a more efficient selection on the X chromosome (Charlesworth et al, 1987; Mank et al, 2010; Meisel and Connallon, 2013); 2) the X chromosome has either more contributing loci or they may have larger effects. Although it is hard to hypothesize about the distribution (number and location) of the selection targets after only 20 generations, we favor the dominance explanation because it is not apparent why the number of selection targets or their effect sizes should be different between the X chromosome and the autosomes.

## MATERIAL AND METHODS

### Experimental set-up

We used the Oregon-R and Samarkand strains inbred by (Chen et al, 2015), and maintained since then at room temperature. The experiment started with 10 replicates, each with a census size of 1500 flies and a starting frequency of 0.3 for the Oregon-R genotype. The 10 replicates were then maintained in parallel at a constant 29°C temperature in dark conditions for 20 generations before sequencing.

### DNA extraction, library preparation, sequencing

Whole-genome sequence data for the parental Oregon-R and Samarkand strains are available from Chen et al. (2015). The 10 evolved replicates in generation F20 were sequenced using Pool-Seq: genomic DNA was prepared after pooling and homogenizing all available individuals of a given replicate in extraction buffer, followed by a standard high-salt extraction protocol (Miller et al. 1988). Barcoded libraries with a mean insert size of 480 bp were prepared using the NEBNext Ultra II DNA Library Prep Kit (E7645L, New England Biolabs, Ipswich, MA) and sequenced on a HiSeq 2500 using a 2 x 125 bp paired-end protocol.

### Establishment of a parental SNPs catalogue

After quality control with FastQC (http://www.bioinformatics.babraham.ac.uk/projects/fastqc/), the raw reads have been demultiplexed and trimmed using ReadTools (Gómez-Sánchez and Schlötterer 2018; version 1.5.2; --mottQualityThreshold 18, --minReadLength 50, --disable5pTrim true). The processed paired-end rends were mapped using NovoAlign (http://novocraft.com; version 3.09; -i 250,75 -F STDFQ -r RANDOM) on the combined *D. melanogaster* reference genome v6.03 (Thurmond et al, 2019). From the processed BAM files, *i*.*e*. without duplicates (using PICARD MarkDuplicates; http://broadinstitute.github.io/picard/; version 2.21.6; REMOVE_DUPLICATES=true VALIDATION_STRINGENCY=SILENT), quality filtered (using samtools (Li et al. 2009); version 1.10; -b -q 20 -f 0×002 -F 0×004 -F 0×008) and re-headed, multi-sample variants calling was done with Freebayes (Garrison and Marth, 2012; version 1.3.1; --use-best-n-alleles 4 --min-alternate-count 3 --ploidy 2 --pooled-continuous --pooled-discrete; version 1.332). Bi-allelic SNPs in regions outside repeats (identified by RepeatMasker, http://www.repeatmasker.org) were extracted from the raw VCF file (Danecek et al, 2011) and filtered using a QUAL value of 1,000 and the 99^th^ percentile averaged coverage as thresholds, leading to a total of 912,289 processed SNPs. A parental SNP was defined as (nearly) fixed difference between parents with a 0/0 (1/1) genotype in the Samarkand parent and 1/1 (0/0) genotype in the Oregon-R parent at the marker position, conditioning for a frequency of the alternate allele lower than 0.01 (if 0/0) or higher than 0.99 (if 1/1). We obtained a final list of 360,517 and 59,280 SNPs on the autosomes and the X chromosome, respectively, equivalent to 1 SNP every 302 bp (397 bp). The frequency of these alleles was measured at F20 (see Table SI 2 for a detailed count of markers at each filtering step) by extracting the number of reads supporting the alternate and reference allele using bcftools (query –H -f ‘%CHROM %POS %REF %ALT %QUAL[%DP][%AO][%RO][%GT]\n’; version 1.9; Li, 2011; piped with sed). The subsequent analyses have been performed with R (version 3.5.0; R Core Team, 2018) and most panels have been done with the ggplot2 R package (Wickham, 2016). For the parental strains, we used the frequency of inversion-diagnostic SNPs to check the inversion status of common cosmopolitan inversions as inversions would impede recombination (Kapun et al, 2014). Both parental strains are homosequential (Fig SI 1). We also checked the density of heterozygous SNPs per parent prior to QUAL filtering (Fig SI 2, top). Both parental strains harbor similar levels of residual variation (Fig SI 2, bottom, bootstrapped Kolmogorov-Smirnov test from Matching R package (Sekhon, 2011) on parental heterozygosity levels; D=0.02, p=0.25).

### Allele frequency tracking

At each SNP we obtained counts for both parental alleles from the VCF file. We polarized allele frequency (AF) for the Oregon-R allele. The frequency of the Samarkand allele is obtained by subtracting the Oregon-R AF from 1. The allele frequency change (AFC) of a given marker is signed; if the Oregon-R AF at F20 is higher (lower) than 30%, the Oregon-R (Samarkand) allele increased in frequency and the AFC is positive (negative). The genome was partitioned in 1,682 non-overlapping genomic windows of 250 parental SNPs (1,444 on the autosomes, 238 on the X chromosome), spanning on average 75 kb (97 kb on the X) where the AF per window was summarized as the median over 250 SNPs. A window position *i* is defined by its center ((right bound-left bound)/2). Markers along the genome are positioned in cM unit, to adjust for heterogeneity in recombination rate along the chromosome. The recombination map of Comeron et al, 2012 was updated to version 6 of the reference genome using the Flybase online Converter (accessed in July 2020). Physical chromosome positions were converted to genetic positions by interpolation (DOQTL R package, Gatti et al, 2014) to avoid SNPs located in the same recombination rate interval to overlap at the cM scale (cf Marey map in Fig SI 3, Mb unit in Fig SI 4). The effective population size, *N*_*e*_, was estimated per replicate for the autosomes and X separately using the poolSeq::estimateNe R function (Taus et al, 2017) from 10,000 randomly picked SNPs and summarized as the median over 1,000 trials, similarly as in Vlachos et al, 2019 (Table SI 3).

### Quantification of the response

For each replicate, we reported the median AF of the Oregon-R allele in each window. We also reported the median coefficient of variation (CV) per chromosome to quantify the deviation around the average AF value per window. We additionally computed the autocorrelation (ACF) in AF between windows using the acf R function. ACF at a given step *k* is defined as the correlation between windows at positions *i* and *i+k*, where *k* is called the lag. We used the distance in Mb at which a significant decrease in ACF was noted (α=5%, below 1.96/√*m, m* the number of windows) as a rough proxy for linkage disequilibrium (LD). We performed neutral simulations mimicking our empirical design (starting frequency of 0.3 for the Oregon-R alleles, 10 replicates, 20 generations, unbiased sex-ratio, census size of 1,500 flies) using MimicrEE2 (Vlachos and Kofler, 2018). From the simulated sync files, we then drew the coverage per SNP from a Poisson distribution (mean=125 reads, estimated from the empirical reads counts) and performed binomial sampling with the sample size equal to the coverage as suggested in Taus et al, 2017, to reproduce Pool-seq sampling noise. To contrast our empirical results with neutral expectations, we computed the pairwise ρ Spearman correlation matrix between all neutral and empirical replicates (10 replicates times 2) per arm (3 major chromosomes), leading to a 10×2×3 entry-matrix. The ρ values were converted to distances (2√(1-ρ)) prior to projecting the distance matrix in two dimensions with Multi-Dimensional Scaling (MDS; Gower, 1966). The significance of the pairwise correlations was assessed with t-tests separately for empirical and neutral replicates, where p-values were adjusted with a Benjamini-Hochberg correction. We performed a sign test for the median AFC to test if the median AFC per major chromosome is higher than 0, where p-values were adjusted with a Benjamini-Hochberg correction.

### Comparisons to other datasets

We qualitatively contrasted our study with two additional E&R studies (Table 1) that are similar in terms of duration and lack inversions but start with hundreds of founder genotypes, and thus heterogeneous starting allele frequencies. To compare studies, we computed the absolute selection coefficient per SNP in one randomly picked replicate; replicate 2 in this study, replicate C from Kelly and Hughes, 2019 (between F0 and F15) and replicate 8 from Barghi et al, 2019 (between F0 and F20) using the same number of SNPs for each study; 248,886 (42,386) sampled SNPs for the autosomes (X). The selection coefficient *s* of each SNP was estimated using the poolSeq::estimateSH R function (Taus et al, 2017) from pseudo-counts; we subtracted (added) a pseudo-count of 1 to fixed (lost) SNPs, as Vlachos et al, 2019. *N*_*e*_ was estimated as described above. We eventually performed pairwise bilateral Wilcoxon-tests between the three *s* distributions for the autosomes and X, where p-values were adjusted with a Benjamini-Hochberg correction.

## ACKNOWLEDGEMENT

We thank all members of the Institut für Populationsgenetik, in particular Anna Maria Langmüller, Neda Barghi and Andreas Futschik for fruitful discussions. We thank Thapasya Vijayan for some preliminary analyses. This work was supported by the Austrian Science Funds (FWF, grant numbers W1225, P29133), and by the European Research Council (ERC) grant “ArchAdapt”. Illumina sequencing was performed at the VBCF NGS Unit (www.viennabiocenter.org/facilities).

## AUTHORS CONTRIBUTIONS

CS designed the experiment. VN performed experiments, DNA extractions and library preparations. VN, MD, CB performed the bioinformatics analysis. CB performed the statistical analysis. VN, MD, CB, CS provided feedback. CB, CS wrote the manuscript with contributions from VN and MD. All authors approved the final manuscript.

## DATA AND SCRIPTS AVAILABILITY

All scripts and data of this study will be available upon publication. Sequence data will be deposited at the European Nucleotide Archive (ENA) under the accession number XXX. Population sync files and scripts will be deposited on Dryad Digital Repository XXX.

