## Supplementary Material for "Highly parallel genomic selection response in replicated *Drosophila melanogaster* populations with reduced genetic variation"

**SUPPLEMENTARY INFORMATION**

**Table SI 1.** Establishment of the parental markers catalogue splitted per arm.

| # SNPs | 2L | 2R | 3L | 3R | 4 | X |
| --- | --- | --- | --- | --- | --- | --- |
| raw | 380,738 | 345,400 | 377,015 | 390,315 | 6,431 | 290,162 |
| after soft filter | 372,822 | 308,627 | 350,985 | 356,763 | 3,328 | 250,067 |
| after hard filters | 228,623 | 155,268 | 211,915 | 212,362 | 520 | 100,793 |
| parental markers | 89,400 | 80,094 | 98,247 | 93,869 | 63 | 59,837 |

**SI Table 2.** *N_e_* estimates per replicate for the autosomes and X separately.

| **Sample** | ***N*_e_ estimate**  **Autosomes / X** |
| --- | --- |
| R1 | 57 / 21 |
| R2 | 55 / 14 |
| R3 | 51 / 22 |
| R4 | 53 / 19 |
| R5 | 55 / 18 |
| R6 | 49 / 22 |
| R7 | 47 / 25 |
| R8 | 59 / 20 |
| R9 | 58 / 17 |
| R10 | 59 / 23 |

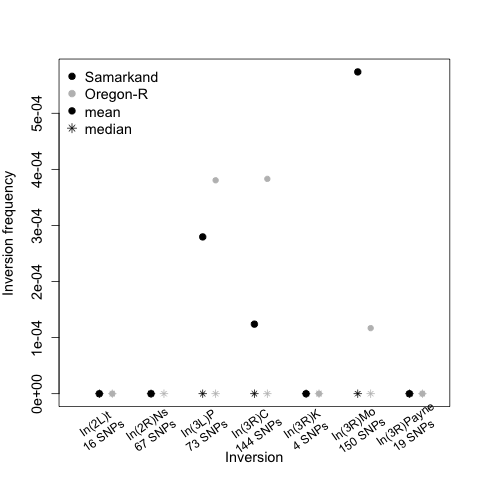

**Figure SI 1.** Inversion status in Samarkand (black) and Oregon-R (gray) parents of known inversions from Kapun et al, 2014.

The mean and median frequencies of the inversions as well as the number of marker SNPs are reported.

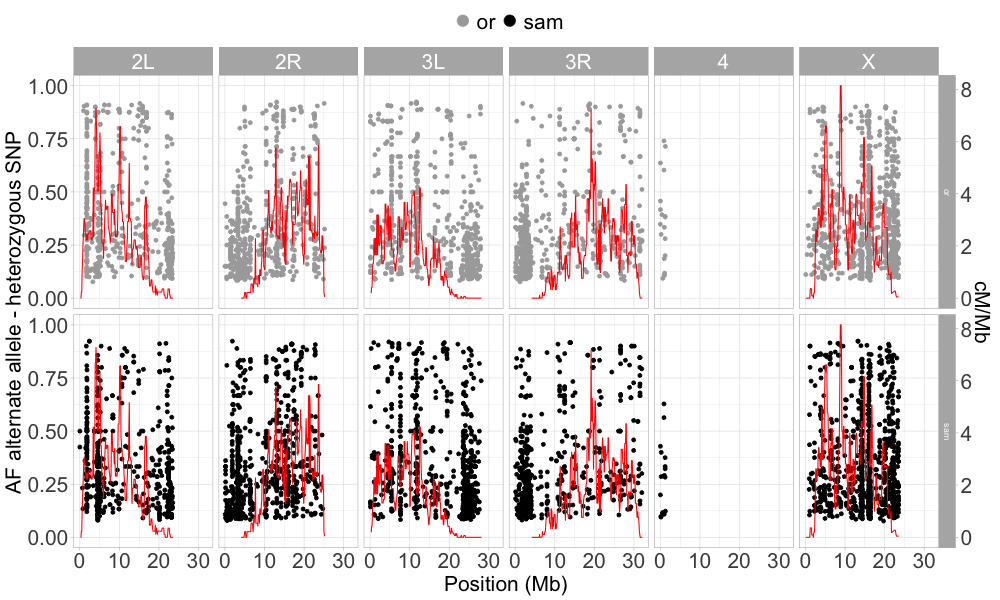

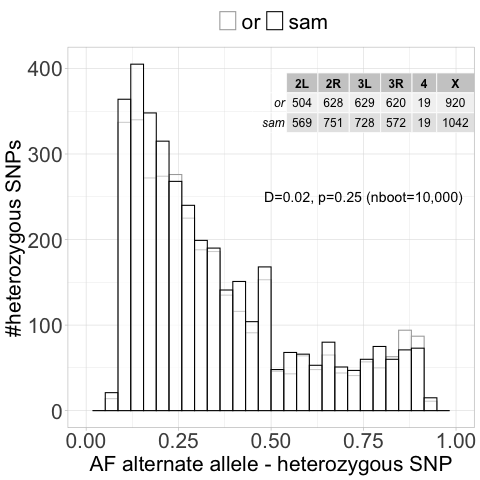

**Figure SI 2.** Residual heterozygosity in Samarkand (gray) and Oregon-R (black).

*Top.* Frequency of the alternate (non-reference) allele at DP-filtered but not QUAL-filtered heterozygous sites per chromosomal location in Samarkand and Oregon-R parental strains as well as recombination rate. *Bottom.* Histograms of the previously plotted frequency per parent. The number of sites are indicated per arm in the inset table. The p-value is obtained with a bootstrapped version of the Kolmogorov-Smirnov test to deal with the presence of ties (n_boot_=10,000).

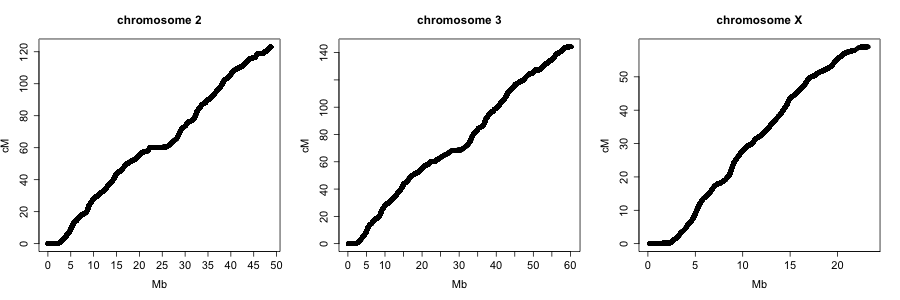

**Figure SI 3.** Marey maps for the major arms showing the genetic position (y-axis) versus the physical position (x-axis) per parental maker. We clearly see the different recombination domains. We did not represent chromosome 4, lacking crossovers (discussed in Hartmann and Sekelsky, 2017).

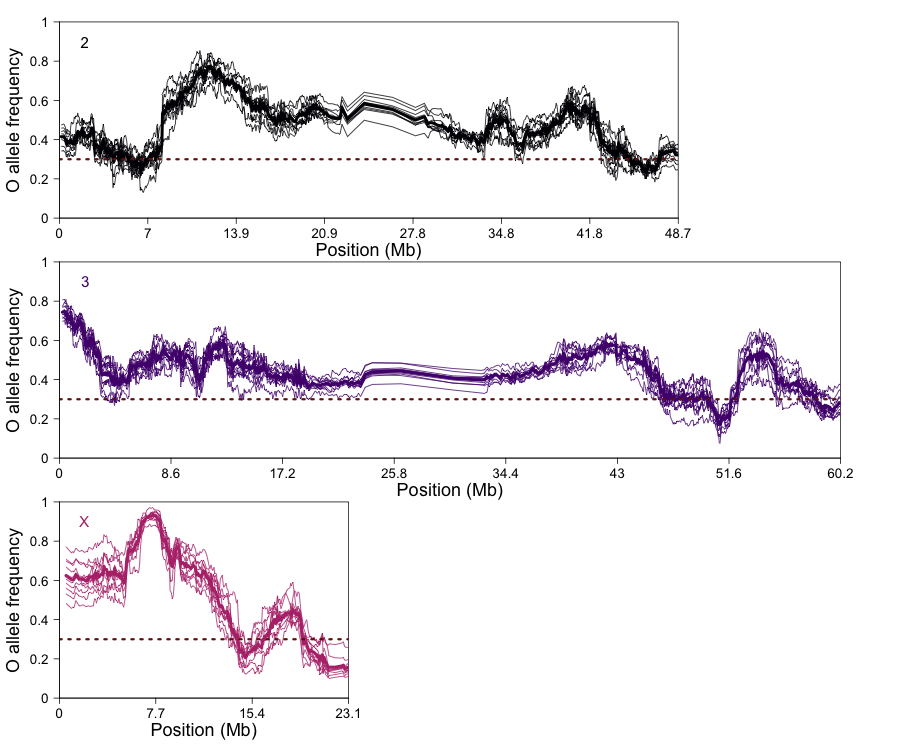

**Figure SI 4.** From cM to Mb unit. Similar Figure as Fig 1. A) but using the Mb unit on the x-axis.
